# Genetic insights into the causal relationship between physical activity and cognitive functioning

**DOI:** 10.1101/2020.10.16.342675

**Authors:** Boris Cheval, Liza Darrous, Karmel W. Choi, Yann C. Klimentidis, David A. Raichlen, Gene E. Alexander, Stéphane Cullati, Zoltán Kutalik, Matthieu P. Boisgontier

## Abstract

Physical activity and cognitive functioning are strongly intertwined. However, the causal relationships underlying this association are still unclear. Physical activity can enhance brain functions, but healthy cognition may also promote engagement in physical activity. Here, we assessed the bidirectional relationships between physical activity and general cognitive functioning using Latent Heritable Confounder Mendelian Randomization (LHC-MR). Association data were drawn from two large-scale genome-wide association studies (UK Biobank and COGENT) on accelerometer-measured moderate, vigorous, and average physical activity (N = 91,084) and cognitive functioning (N = 257,841). After Bonferroni correction, we observed significant LHC-MR associations suggesting that increased fraction of both moderate (b = 0.32, CI_95%_ = [0.17,0.47], P = 2.89e-05) and vigorous physical activity (b = 0.22, CI_95%_ = [0.06,0.37], P = 0.007) lead to increased cognitive functioning. In contrast, we found no evidence of a causal effect of average physical activity on cognitive functioning, and no evidence of a reverse causal effect (cognitive functioning on any physical activity measures). These findings provide new evidence supporting a beneficial role of moderate and vigorous physical activity (MVPA) on cognitive functioning.

## Introduction

Multiple cross-sectional and longitudinal studies have shown that physical activity and cognitive functioning are strongly intertwined and decline through the course of life (Cheval, Orsholits, et al., 2020; Cheval, Sieber, et al., 2018; DiPietro, 2001; Levy, 1994; Sebastiani et al., 2020). However, the evidence of causality of this relationship remains unclear Previous results have shown that physical activity can improve cognitive functioning (Angevaren et al., 2008; Baumgart et al., 2015; Blondell et al., 2014; Colcombe & Kramer, 2003; Hamer et al., 2018; Morgan et al., 2012; Sofi et al., 2011), but recent studies have also suggested that well-functioning cognitive skills can influence engagement in physical activity (Cheval, Bacelar, et al., 2020; Cheval et al., 2022; Cheval, Orsholits, et al., 2020; Cheval et al., 2019; Daly et al., 2015; Lindwall et al., 2012; Sabia et al., 2017; Snowden et al., 2011; Young et al., 2015).

Several mechanisms could explain how physical activity enhances general cognitive functioning (Colcombe & Kramer, 2003; Colzato et al., 2018; Cotman & Berchtold, 2002; Cotman et al., 2007; Hillman et al., 2008; Lisanne et al., 2018; Raichlen & Alexander, 2017; Roig et al., 2013). For example, physical activity can increase brain plasticity, angiogenesis, synaptogenesis, and neurogenesis primarily through the upregulation of growth factors (e.g., brain-derived neurotrophic factor; BDNF) (Cotman & Berchtold, 2002; Cotman et al., 2007; Hillman et al., 2008). In addition, the repetitive activation of higher-order brain functions (e.g., planning, inhibition, and reasoning) required to engage in physical activity may contribute to the improvement of these functions (Frith & Loprinzi, 2018; Raichlen & Alexander, 2017). In turn, other mechanisms could explain how cognitive functioning may affect physical activity. For example, cognitive functioning may be required to counteract the innate attraction to effort minimization and thereby influence a person’s ability to engage in physically active behavior (Cheval, Daou, et al., 2020; Cheval, Radel, et al., 2018; Cheval et al., 2019; Cheval, Tipura, et al., 2018). Of note, these mechanisms are not mutually exclusive and could therefore lead to bidirectionally reinforcing relationships (i.e., positive feedback loop) between physical activity and cognitive functioning (Choi et al., 2019). Thus, there is a mechanistic explanation theoretically supporting the associations between physical activity and cognitive function.

Although these studies point to a potential mutually beneficial interplay between physical activity and cognitive functioning across the lifespan, these findings mainly stem from observational designs and analytical methods that cannot fully rule out the influence of social, behavioral, and genetic confounders (Choi et al., 2019). While randomized controlled trials minimizing these potential counfounds have been conducted (Northey et al., 2018), they were typically based on small sample sizes (n<100) that can bias the estimations (Northey et al., 2018). Critically, these trials only investigated the effect of physical activity on cognitive functioning, not the opposite. Accordingly, current evidence on the causal association between physical activity and cognitive functioning and on whether this association is one or two-way could be considered weak. Because Mendelian Randomization (MR) is less vulnerable to confounding or reverse causation than conventional approaches in observational studies (Byrne et al., 2017; Davies, Holmes, et al., 2018), this method is particularly appropriate to address this knowledge gap.

MR is an epidemiological method in which the randomized inheritance of genetic variation is considered as a natural experiment to estimate the potential causal effect of a modifiable risk factor (exposure) on health-related outcomes in an observational design (Byrne et al., 2017; Davies, Holmes, et al., 2018). MR draws on the assumption that genetic variants are associated with the exposure, because they are randomly allocated at conception, are less associated with other risk factors that may be confounders of the exposure and the outcome, and are immune to reverse causality since diseases or health-related outcomes have no reverse effect on genetic variants. Accordingly, if an exposure (e.g., physical activity) causally affects an outcome (e.g., cognitive function), the genetic variants that influences this exposure is expected to affect the outcome to a proporitional degree if no separate pathway exists by which these genetic variants can affect the outcome (Choi et al., 2019). In other words, genetic variants associated with an exposure of interest can serve as instruments (or proxies) for estimating the causal association with an outcome (see Figure 1 for the conceptual illusation of the MR method).

**Figure 1.**
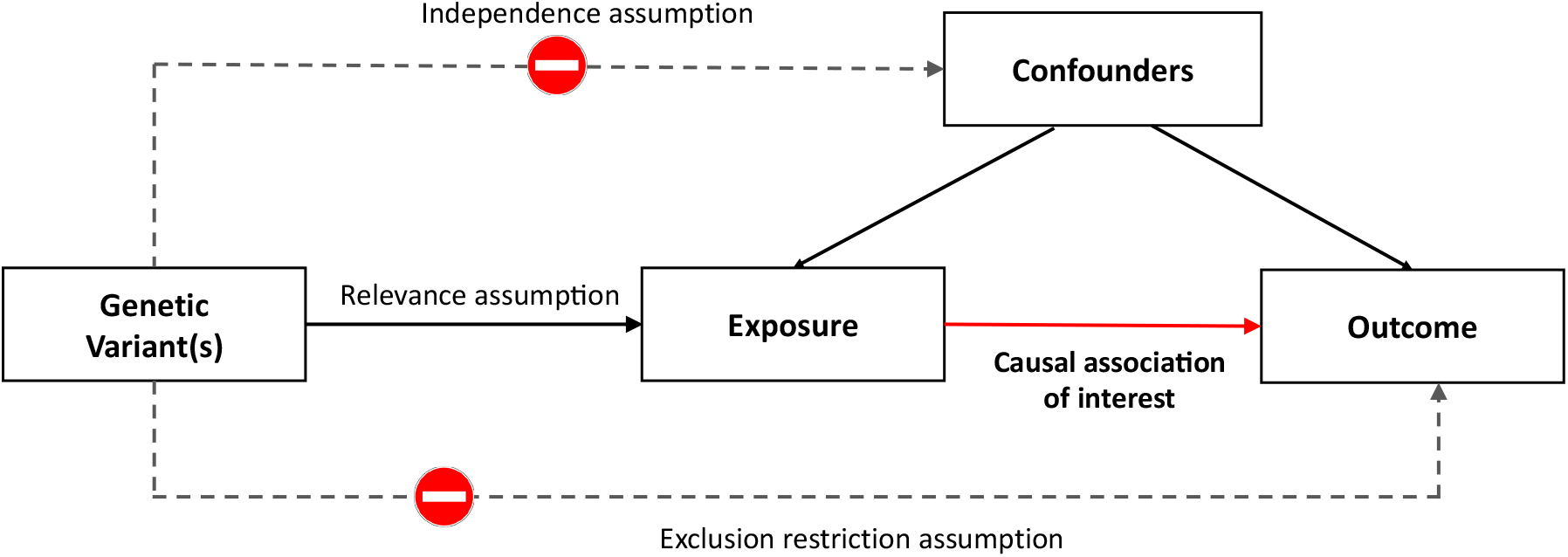
Conceptual illustration of the MR method. Notes. The causal association of interest is between the exposure (e.g., physical activity) and the outcome (e.g., cognitive function). Relevance assumption states that the genetic instruments are strongly associated with the exposure but are not associated with the counfouders. The exclusion restriction assumption states that the genetic instruments are only indirectly associated with the outcome via the exposure. Thus, the solid paths are expected to exist, while the dashed paths are expected to be nonsngificant according to the core MR assumptions.

We used a newly-developed MR method showing improved power to simultaneously estimate the bidirectional causal effects between physical activity and cognitive functioning (Darrous et al., 2021). In a two-sample MR design, genetic instruments can be obtained from summary statistics of nonoverlapping large-scale genome-wide association studies (GWAS). That is, the genetic instruments for the exposure and the genetic instruments for the outcome can be obtained from separate studies (Hemani et al., 2018). This is an outstanding advantage for estimating the causal relationships between two traits (e.g., cognitive functioning and physical activity) because a trait does not necessarily need to be assessed in both samples (Hemani et al., 2018). Here, the causal estimates were modelled based on recently available summary statistics from large-scale GWAS of accelerometer-measured physical activity (Klimentidis et al., 2018), and general cognitive functioning (Davies, Lam, et al., 2018; Lee et al., 2018). Since it has been suggested that the intensity of physical activity can be an important consideration (Szuhany et al., 2015), we assessed whether the causal effect estimates on cognitive functioning were dependent on physical activity intensity (i.e., moderate vs. vigorous vs. average intensity). Of note, as existiting literature suggests reciprocal associations between physical activity and cognitive function, we applied bidirectional MR to examine the causal link from physical activity to cognitive function and from cognitive function to physical activity.

## Methods

### Data sources and instruments

This study used de-identified GWAS summary statistics from original studies that were approved by relevant ethics committees. The current study was approved by the Ethics Committee of Geneva Canton, Switzerland (CCER-2019-00065). The available summary-level data were based on 257,841 samples for general cognitive functioning and 91,084 samples for accelerometer-based physical activity. Participants’ age ranged from 40 to 69 years in the UK Biobank and from 8 to 96 years in the COGENT consortium.

### Physical activity

*Accelerometer-measured physical activity* was assessed based on summary statistics from a recent GWAS (Klimentidis et al., 2018), analyzing accelerometer-based physical activity data from the UK Biobank. In the UK Biobank, about 100,000 participants wore a wrist-worn triaxial accelerometer (Axivity AX3) that was set up to record data for seven days. Individuals with less than 3 days (72 h) of data or not having data in each 1-hour period of the 24-h cycle or for whom the accelerometer could not be calibrated were excluded. Data for non-wear segments, defined as consecutive stationary episodes ≥ 60 min where all three axes had a standard deviation < 13 mg, were imputed. The details of data collection and processing can be found elsewhere (Doherty et al., 2017). We examined three measures derived from the three to seven days of accelerometer wear: the average acceleration in milli-gravities (mg) that includes acceleration > 100 mg, the fraction of accelerations > 100 mg and < 425 mg to estimate moderate physical activity, and fraction of accelerations ≥ 425 mg to estimate vigorous physical activity (Klimentidis et al., 2018). As previous reported (Hildebrand et al., 2014), 425 mg cut-off was chosen because it corresponds to vigorous intensity (6 METS). The GWAS for average physical activity (*n*_*max*_ = 91,084) identified 2 independent genome-wide significant SNPs (P < 5e-09), with a SNP-based heritability of ∼ 14%.

As for the other two physical activity measures, the fractions of accelerations corresponding to moderate and vigorous physical activity were obtained by running new GWAS on the decomposed acceleration data from UK Biobank using the BGENIE software (Bycroft et al., 2018). The phenotype for moderate physical activity was limited to acceleration magnitudes ranging from 100 to < 425 mg, whereas vigorous physical activity was limited to acceleration magnitudes ranging from 425 to 2,000 mg. These acceleration fractions were adjusted for age, sex, and the first 40 principal component (PC), and the analyzed individuals were restricted to unrelated white-British. The two datasets of average physical activity summary statistics, alongside the moderate and vigorous physical activity summary statistics, were used in Latent Heritable Confounder Mendelian Randomization (LHC-MR) to investigate the possible bidirectional effect that exists between these physical activity traits and cognitive functioning.

### General cognitive functioning

*General cognitive functioning* was assessed based on summary statistics from a recent GWAS combining cognitive and genetic data from the UK Biobank and the COGENT consortium (N = 257,841) (Lee et al., 2018). The phenotypes of these cohorts are well-suited to meta-analysis because their pairwise genetic correlation has been shown to be high (Davies, Lam, et al., 2018). In the UK Biobank (*n*_*max*_ = 222,543) participants were asked to complete 13 multiple-choice questions that assessed verbal and numerical reasoning. For verbal reasoning, a typical question was “bud is to flower what child is to …?”, and possible answers presented to the participants are “Grow”, “Develop”, “Improve”, “Adult”, or “Old”. For numerical reasoning, a typical question was “150…137…125…114…104… what comes next?” with possible answers being “96”, “95”, “94”, “93”, or “92” (Lee et al., 2018). The verbal and numerical reasoning score was based on the number of questions answered correctly within a two-minute time limit. Each respondent took the test up to four times. This test was designed as a measure of fluid intelligence. The phenotype consists of the mean of the standardized score across the measurement occasions for a given participant. In the COGENT consortium (*n*_*max*_ = 35,298), general cognitive function is statistically derived from a principal components analysis of individual scores on a neuropsychological test battery, such as the Verbal or spatial N-Back working memory task, Stroop Test, the Trail Making Test, or the Wechsler Adult Intelligence Scale (Trampush et al., 2017). Details on the test battery are available in the supplementary material of Davies et al. (2016). Of note, Davies et al. (2016) demonstrated that two general cognitive function components extracted from different sets of cognitive tests on the same participants exhibit a high correlation, addressing the fact that different cohorts relied on different cognitive tests. Thus, the phenotype estimates overall cognitive functioning and is relatively invariant to the battery used and specific cognitive abilities assessed (Johnson et al., 2008; Panizzon et al., 2014). These COGENT data used to assess general cognitive functioning were also used in another GWAS study (Davies, Lam, et al., 2018). The GWAS identified 226 independent genome-wide significant SNPs, with a SNP-based heritability of ∼20%.

### Statistical analysis

MR is a statistical approach for causal inference that can overcome the weaknesses of traditional observational studies (Byrne et al., 2017; Davies, Holmes, et al., 2018). MR-based effect estimates rely on three main assumptions (Lawlor et al., 2008), stating that genetic instruments i) are strongly associated with the exposure (relevance assumption), ii) are independent of confounding factors of the exposure-outcome relationship (independence assumption), and iii) are not associated to the outcome conditional on the exposure and potential confounders (exclusion restriction assumption). Well-powered GWAS offer multiple genetic instruments that are strongly associated with exposures of interest (cognitive functioning or physical activity in our case), which validates the relevance assumption. Each of these genetic variants (instruments) provides a causal effect estimate of the exposure on the outcome, which can be in turn combined through meta-analysis using inverse-variance weighting (IVW) to obtain an overall estimate. The second and third assumptions are less easily validated and can be violated in the case of a heritable confounder affecting the exposure-outcome relationship and biasing the causal estimate. Such confounders can give rise to instruments with proportional effects on the exposure and outcome, hence violating the INstrument Strength Independent of Direct Effect (InSIDE) assumption requiring the independence of the exposure and direct outcome effects. There have been several extensions to the common IVW method of MR analysis, including MR-Egger, which allows for directional pleiotropy of the instruments and attempts to correct the causal regression estimate. Other extensions, such as median and mode-based estimators, assume that at least half of or the most “frequent” genetic instruments are valid/non-pleiotropic. However, despite these extensions and relaxed assumptions, all these classical MR methods are notably underpowered^32^ and still suffer from two major limitations. First, they only use a subset of markers as instruments (genome-wide significant markers), which often dilutes the true relationship between traits. Second, they ignore the presence of a potential latent heritable confounder of the exposure-outcome relationship (e.g., body mass index, educational attainement, level of physical activity at work, or material deprivation).

LHC-MR also uses GWAS summary statistics (Darrous et al., 2021), but importantly, this new method appropriately uses genome-wide genetic markers to estimate bidirectional causal effects, direct heritability, and confounder effects while accounting for sample overlap. LHC-MR can be viewed as an extension of the linkage disequilibrium score regression (LDSC) (Bulik-Sullivan et al., 2015), designed to estimate trait heritability, in that it models all genetic marker effects as random, but additionally estimates bidirectional causal effect, as well as other parameters. LHC-MR extends the standard two-sample MR by modeling a latent (unmeasured) heritable confounder that has an effect on the exposure and outcome traits. This allows LHC-MR to differentiate SNPs based on their co-association to a pair of traits and distinguish heritable confounding that leads to genetic correlation from actual causation. Thus, the unbiased bidirectional causal effect between these two traits are estimated simultaneously along with the confounder effect on each trait (Figure 2a-b). The LHC-MR framework, with its multiple pathways through which SNPs can have an effect on the traits, as well as its allowance for null effects, make LHC-MR more precise at estimating causal effects compared to standard MR methods (i.e., MR egger, weighted median, inverse variance weigthed, simple mode, and weighted mode).

**Figure 2.**
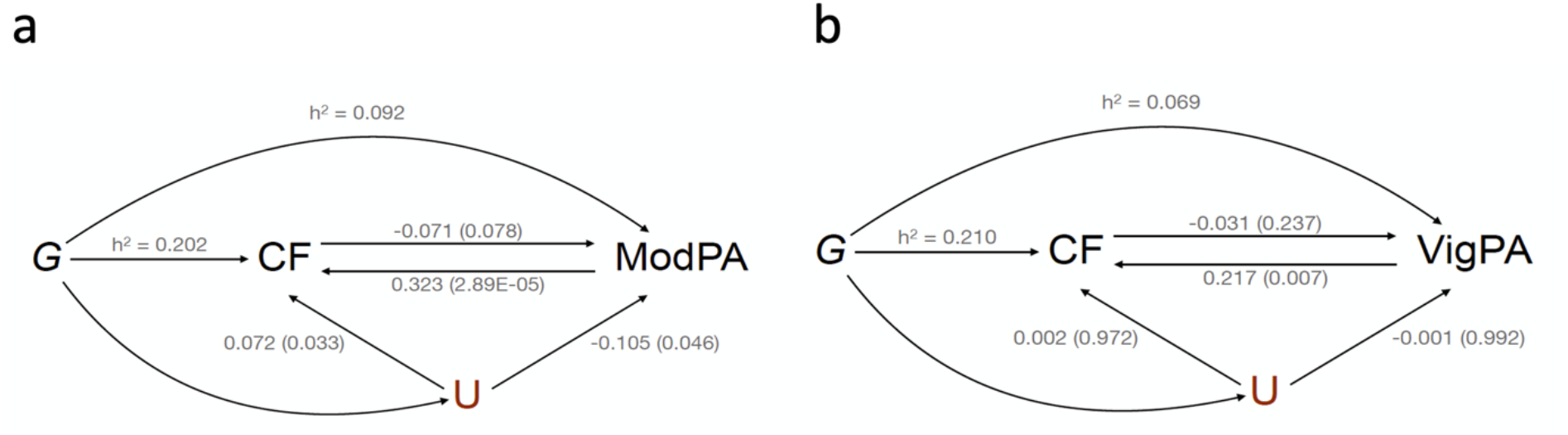
Visual representation of the model in LHC-MR. Notes. G = Genetic instruments; CF = general cognitive functioning; a. For moderate physical activity (ModPA); b. For vigorous physical activity (VigPA); U = Latent heritable confounder; h^2^ = direct heritability. Each figure includes the bidirectional causal effects between the two traits as well as the confounder effects on each of them. Coefficients are beta values. P-values are indicated in brackets. The models were adjusted for age, sex, genotyping chip, first ten genomic principal components (PC), center, and season (month) of wearing accelerometer.

The likelihood function for LHC-MR, which is derived from the mixture of different pathways through which the genome-wide SNPs can have an effect (acting on either the exposure, the outcome, the confounder, or the combinations of these three), is then optimized given random starting values for the parameters it can estimate. The optimization of the likelihood function then yields the maximum likelihood estimate (MLE) value for a set of estimated parameters, including the bidirectional causal effect between the exposure and the outcome as well as the strength of the confounder effect on each of those two traits. The standard errors of each of the parameters estimated using LHC-MR were obtained by implementing a block jackknife procedure where the SNP effects are split into blocks, and the MLE is computed again in a leave-one-block-out fashion. The variance of the estimates can then be computed from the results of the various MLE optimizations. Furthermore, the causal estimates obtained from LHC-MR are on the scale of 1 standard deviation (SD) outcome difference upon a 1 SD exposure change due to the use of standardized summary statistics for the two traits.

A sensitivity analysis in which the model was further adjusted for baseline self-reported level of physical activity at work, walking or standing at work, and the Townsend Deprivation Index was conducted.

## Results

Three measures derived from accelerometer wear were used as a proxy for physical activity: average, moderate, and vigorous physical activity. These three measures were used in LHC-MR to investigate the possible bidirectional causal effects between them and cognitive functioning. The model tested was adjusted for age, sex, genotyping chip, first ten genomic principal components (PC), center, and season (month) of wearing accelerometer. The Bonferroni correction was used to control for familywise error rates, yielding an α = 0.05 / (2 directions × 3 tests) = 0.008.

### Average physical activity and general cognitive functioning

LHC-MR applied to summary statistics belonging to model 1 showed no evidence for a potential causal effect of average physical activity on cognitive functioning (b = 0.245, CI_95%_ = [-0.01,0.50], P = 0.065) (Table 1, Figure 3) and no evidence for the reverse causal effect (b = -0.145, CI_95%._ = [-0.26,-0.03], P = 0.013 [α = 0.008]). Similarly, standard MR methods such as IVW, MR Egger, weighted median, simple mode, and weighted mode yielded non-significant causal estimates in either direction (Table 2), using 129 genome-wide significant single nucleotide polymorphisms (SNPs) as instruments for cognitive functioning and 6 SNPs for average acceleration.

**Table 1.** Latent Heritable Confounder Mendelian Randomization (LHC-MR) results for the association between accelerometer-measured physical activity and general cognitive functioning.

| Parameter | Cognitive Functioning |  | Physical Activity |  | Cognitive Functioning<br>→ Physical Activity | Physical Activity<br>→ Cognitive Functioning |
| --- | --- | --- | --- | --- | --- | --- |
|  | Heritability | t | Heritability | t |  |  |
| Average accelerometer-measured physical activity |  |  |  |  |  |  |
| Estimate | 0.207 | 0.033 | 0.123 | -0.011 | -0.145 | 0.245 |
| P-value | 2.67E-115 | 0.612 | 4.41E-28 | 0.816 | 0.013 | 0.065 |
| Moderate accelerometer-measured physical activity (fraction of acceleration > 100 mg and < 425 mg) |  |  |  |  |  |  |
| Estimate | 0.202 | 0.072 | 0.092 | -0.105 | -0.071 | 0.323 |
| P-value | 1.13E-165 | 0.032 | 5.98E-29 | 0.046 | 0.078 | 2.89e-05 |
| Vigorous accelerometer-measured physical activity (fraction of acceleration ≥ 425 mg) |  |  |  |  |  |  |
| Estimate | 0.210 | 0.002 | 0.069 | -0.001 | -0.031 | 0.212 |
| P-value | 6.75E-157 | 0.972 | 3.95E-25 | 0.992 | 0.237 | 0.007 |

Notes. Parameters estimates and their p-values obtained from the LHC-MR optimized model with the maximum likelihood. Bidirectional associations from cognitive functioning to physical activity and from physical activity to cognitive functioning are reported. t = effect of the confounder. Bonferroni corrected $\alpha = 0.008$ .

**Table 2.** Standard Mendelian Randomization (MR) results for the association between accelerometer-based physical activity and general cognitive functioning.

| Exposure | Outcome | MR method | Valid SNPs | Causal estimate | SE | P-value |
| --- | --- | --- | --- | --- | --- | --- |
| Average accelerometer-based physical activity |  |  |  |  |  |  |
| Cognitive Functioning | Physical Activity | MR Egger | 129 | 0.015 | 0.185 | 0.935 |
|  |  | Weighted median | 129 | -0.027 | 0.036 | 0.440 |
|  |  | Inverse variance weighted | 129 | -0.011 | 0.032 | 0.723 |
|  |  | Simple mode | 129 | -0.102 | 0.116 | 0.376 |
|  |  | Weighted mode | 129 | -0.084 | 0.111 | 0.452 |
| Physical Activity | Cognitive Functioning | MR Egger | 4 | -2.833 | 1.148 | 0.069 |
|  |  | Weighted median | 4 | 0.017 | 0.062 | 0.782 |
|  |  | Inverse variance weighted | 4 | -0.088 | 0.127 | 0.488 |
|  |  | Simple mode | 4 | 0.020 | 0.076 | 0.801 |
|  |  | Weighted mode | 4 | 0.023 | 0.074 | 0.770 |
| Moderate accelerometer-based physical activity (fraction of acceleration > 100 mg and < 425 mg) |  |  |  |  |  |  |
| Cognitive Functioning | Physical Activity | MR Egger | 129 | -0.054 | 0.181 | 0.766 |
|  |  | Weighted median | 129 | -0.032 | 0.037 | 0.389 |
|  |  | Inverse variance weighted | 129 | -0.012 | 0.032 | 0.710 |
|  |  | Simple mode | 129 | -0.059 | 0.106 | 0.575 |
|  |  | Weighted mode | 129 | -0.031 | 0.091 | 0.729 |
| Physical Activity | Cognitive Functioning | MR Egger | 106 | 0.325 | 0.319 | 0.310 |
|  |  | Weighted median | 106 | -0.001 | 0.021 | 0.981 |
|  |  | Inverse variance weighted | 106 | 0.023 | 0.022 | 0.309 |
|  |  | Simple mode | 106 | -0.017 | 0.057 | 0.767 |
|  |  | Weighted mode | 106 | -0.010 | 0.050 | 0.837 |
| Vigorous accelerometer-based physical activity (fraction of acceleration ≥ 425 mg) |  |  |  |  |  |  |
| Cognitive Functioning | Physical Activity | MR Egger | 129 | 0.009 | 0.149 | 0.952 |
|  |  | Weighted median | 129 | 0.018 | 0.036 | 0.623 |
|  |  | Inverse variance weighted | 129 | 0.002 | 0.026 | 0.939 |
|  |  | Simple mode | 129 | 0.021 | 0.097 | 0.829 |
|  |  | Weighted mode | 129 | 0.021 | 0.088 | 0.812 |
| Physical Activity | Cognitive Functioning | MR Egger | 88 | 0.151 | 0.335 | 0.653 |
|  |  | Weighted median | 88 | -0.035 | 0.022 | 0.108 |
|  |  | Inverse variance weighted | 88 | -0.016 | 0.020 | 0.432 |
|  |  | Simple mode | 88 | -0.065 | 0.060 | 0.286 |
|  |  | Weighted mode | 88 | -0.059 | 0.052 | 0.257 |

Notes. Causal estimates from 5 standard Mendelian Randomization (MR) methods on alternating exposure and outcome traits. For both moderate and vigorous physical activity as exposure, the cutoff was decreased to $6.33\text{e-}5$ because of the low number of genome wide significant single nucleotide polymorphisms (SNPs) to use as instruments. Corrected $\alpha = 0.008$ .

**Figure 3.**
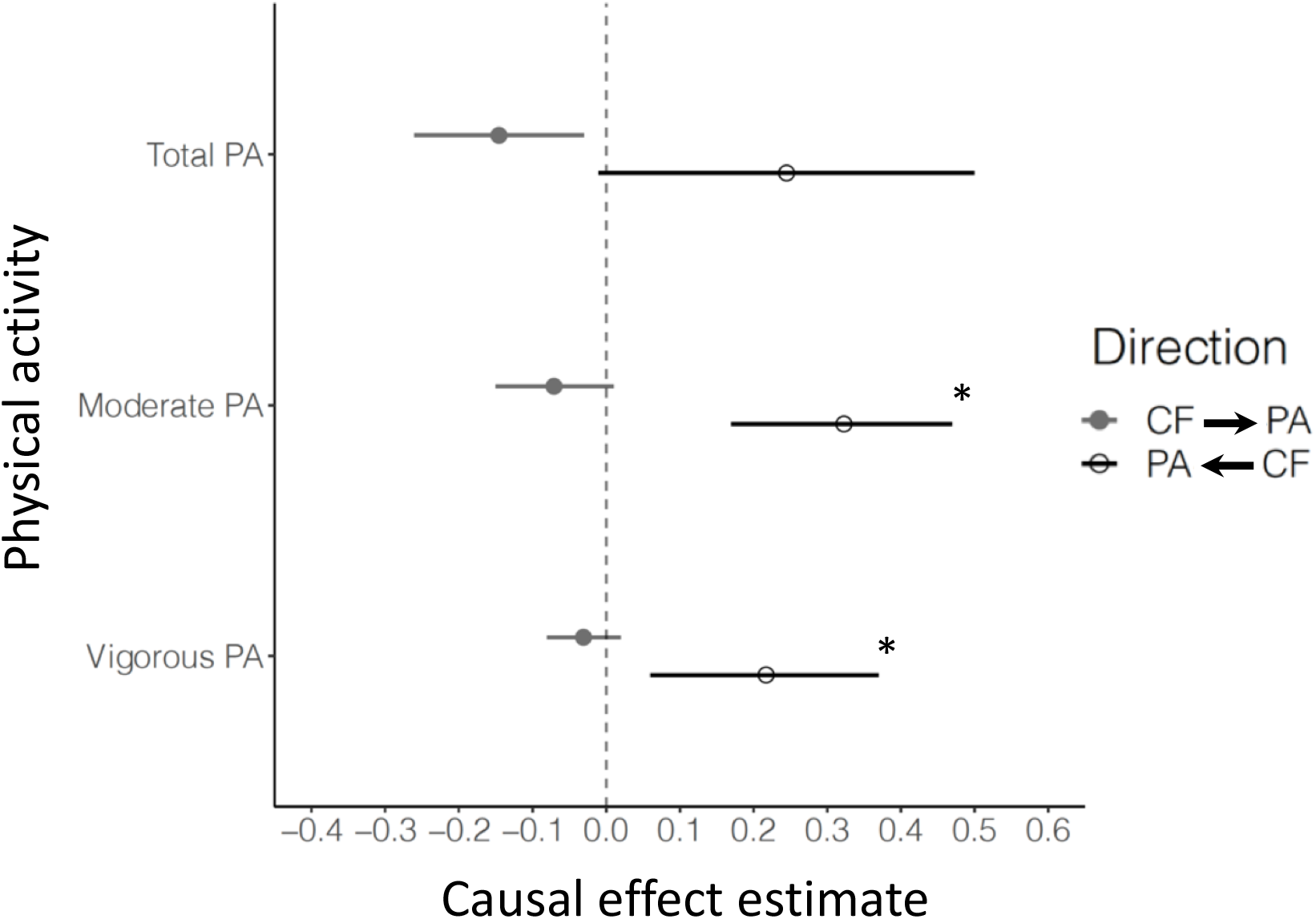
LHC-MR plots for the association between accelerometer-based physical activity and general cognitive functioning. Notes. This modified dot-and-whisker plot reports the causal estimate between general cognitive functioning (CF) as exposure and varying physical activity (PA)-related traits as outcome. The forward (CF → PA) and reverse (PA → CF) causal estimates are shown in two different colors as dots (grey and white) with 95% CI whiskers (grey and black). Average PA = average of overall accelerations. Moderate PA = fraction of acceleration corresponding to moderate physical activity (> 100 mg and < 425 mg). Vigorous PA = fraction of acceleration corresponding to vigorous physical activity (≥ 425 mg). The models were adjusted for age, sex, genotyping chip, first ten genomic principal components (PC), center, and season (month) of wearing accelerometer. * = significant effect after Boneferroni correction (i.e., p-value < .008).

### Moderate physical activity and general cognitive functioning

LHC-MR applied to the fraction of accelerations corresponding to moderate physical activity showed a potential positive causal effect of moderate physical activity on greater cognitive functioning (b = 0.32, CI_95%_ = [0.17,0.47], P = 2.89e-05) (Table 1, Figure 3). We found no evidence for the reverse causal effect (b = -0.071, CI_95%._ = [-0.15, 0.01], P = 0.078 [α = 0.008]). As was found with average physical activity, there was no evidence for the presence of a heritable confounder. Standard MR methods yielded non-significant causal estimates in both directions (Table 2).

### Vigorous physical activity and general cognitive functioning

LHC-MR applied to the fraction of accelerations corresponding to vigorous physical activity on cognitive functioning showed a potential positive causal effect of vigorous physical activity on greater cognitive functioning (b = 0.22, CI_95%_ = [0.06,0.37], P = 0.007) (Table 1, Figure 3). We found no evidence for the reverse causal effect (b = -0.031, CI_95%._ = [-0.08, 0.02], P = 0.237 [α = 0.008]). As was found with average and moderate physical activity, there was no evidence for the presence of a heritable confounder. Of note, the coefficient of this causal effect was qualitatively weaker than of the causal effect of moderate physical activity on cognitive functioning (b = 0.22 vs. b = 0.32). Standard MR methods yielded non-significant causal estimates in both directions (Table 2).

### Sensitivity analyses

We tested another model where an extra adjustment had been done for the baseline self-reported level of physical activity at work, walking or standing at work, and the Townsend Deprivation Index. LHC-MR applied to summary statistics emerging from this second model showed consistent results with that of the first model (b = 0.22, CI_95%_ = [-0.05,0.50], P = 0.111 and b = -0.090, CI_95%_ = [-0.23,0.05], P = 0.200, respectively). Both models showed no evidence for the presence of a heritable confounder. Due to the similarity in results between these models, we did not conducted this second model on moderate and vigoruous physical activity.

## Discussion

### Main findings

This study used a genetically informed method that provides evidence of putative causal relations to investigate the bidirectional associations between accelerometer-based physical activity and general cognitive functioning. Drawing on large-scale GWAS, we found evidence for potential causal effects, suggesting that higher levels of moderate and vigorous physical activity lead to increased cognitive functioning. In the opposite direction, we did not observe evidence of a causal effect of cognitive functioning on physical activity. Hence, our study suggests a favorable effect of moderate and vigorous physical activity on cognitive functioning, but does not provide evidence that increased cognitive functioning promotes engagement in more physical activity.

### Comparison with previous studies

Previous reviews and meta-analyses of observational studies showed a beneficial effect of physical activity on cognitive functioning (Baumgart et al., 2015; Morgan et al., 2012; Raichlen & Alexander, 2017; Sofi et al., 2011). However, the evidence arising from intervention studies was inconclusive (Angevaren et al., 2008; Colcombe & Kramer, 2003; Erickson et al., 2019; Sabia et al., 2017; Snowden et al., 2011; Young et al., 2015). It has been argued that these inconsistencies may primarily be attributed to the design-specific tools used to assess physical activity (Sabia et al., 2017). Specifically, many observational studies rely on self-reported measures of physical activity, whereas intervention studies often rely on accelerometer-measured physical activity, or have people exercising under monitored conditions. In other words, evidence of a favorable effect of physical activity on cognitive functioning may have emerged in observational studies because of the self-reported nature of the measures they typically used. Yet, in our study, results are based on accelerometer-assessed physical activity, thereby partially ruling out this explanation. Therefore, our findings further support the literature that demonstrated a protective role of physical activity on cognitive functioning and extend it by doing so using an accelerometer-based measure. Our findings are in line with recent MR-based results showing a protective effect of objectively assessed, but not self-reported, physical activity on the risk of depression (Choi et al., 2019).

Of note, results obtained from LHC-MR differed from those obtained with standard MR methods. At least three key differences in the methods can explain this divergence: i) standard MR uses only genome-wide significant markers, ii) standard MR is biased in case of sample overlap (as is the case in this study) and hence their estimate may be biased towards the observational correlation, and iii) LHC-MR explicitly models correlated pleiotropy unlike standard MR. Accordingly, our results obtained from LHC-MR are expected to be more robust than those obtained from standard MR. Since LHC-MR could not find evidence for the presence of a heritable confounder, correlated pleiotropy is less likely, or there might be multiple confounders with opposite effects cancelling each other out. This finding highlights that the main reason for the difference between LHC-MR and classical MR methods is statistical power. For testing the reverse causal effect (cognition on physical activity), we had numerous instruments available, ensuring that all MR-methods are well-powered and yielding the same (null effect) conclusion. The forward effect (physical activity on cognition) relied on only a few (weak) instruments, rendering classical MR methods notably underpowered. This is the type of situation in which methods such as LHC-MR, which leverage genome-wide genetic markers, are crucial to facilitate discovery. It is important to point out that while the statistical conclusion from classical and LHC-MR methods differ, their effect estimates are not significantly different, suggesting that there is no discrepancy in the results, but that they have different precision. Finally, we acknowledge that LHC-MR assumptions may be violated and results should thus still be considered cautiously. Yet, while the assumptions of LHC-MR may not hold, the assumptions of the other five methods are known not to hold because of insufficient genome-wide significant instrument.

To the best of our knowledge, our study is the first to investigate the potential causal relationship between physical activity and cognitive functioning using a genetically informed method. We are aware of only two other, non-genetic studies that examined the potential bidirectional associations between physical activity and cognitive functioning (Cheval, Orsholits, et al., 2020; Daly et al., 2015). In contrast to the present study, those two studies observed a positive influence of cognitive functioning on physical activity. At least two factors can explain the differences in the results observed. First, both those studies are based on longitudinal assessment (Granger causality) of the two traits, while our approach is based on a genetically instrumented causal inference technique (LHC-MR). Second, these studies draw on self-reported physical activity rather than accelerometer-measured physical activity, which may not accurately reflect the objective level of physical activity.

Our results obtained with recently-improved genetically-informed analyses (LHC-MR) highlight the potential critical role of physical activity, specifically of moderate and vigorous intensity, on cognitive functioning. However, it should be noted that the estimated effect of moderate physical activity on cognitive functioning was about 1.5 times stronger in magnitude than the effect of vigorous physical activity. To the best of our knowledge, this study is the first to assess and compare the causal relationships of moderate and vigorous physical activity with cognitive functioning using a genetically-informed method based on large-scale datasets. Alhough additional evidence are needed, this study confirms the importance to examine the extent to which the intensity of physical activity moderates the effects observed on cognitive functioning (Szuhany et al., 2015).

The LHC-MR method revealed two causal relations that are consistent with each other. Importantly, these findings are consistent with theoretical and experimental work explaining the mechanisms underlying the association between the physical activity and cognitive functioning. Results obtained with both the LHC and standard MR methods showed no evidence of an effect of average physical activity on cognitive functioning. This finding can likely be explained by physical activities of low intensity (i.e., < 100 mg) that are part of the average physical activity, which further suggests that physical activity should be of moderate-to-vigorous intensity to benefit cognitive functioning.

The absence of evidence for a reverse causal effect of cognitive functioning on physical activity may be partly explained by the lower power of this analysis due to smaller sample size of the GWAS of physical activity (n = 91,084) compared to the sample size of the GWAS of cognitive functioning (n = 257,841). This absence of evidence contrasts with other studies arguing that cognitive functioning is critical for supporting engagement in physical activity (Cheval, Radel, et al., 2018; Cheval et al., 2019; Cheval, Tipura, et al., 2018). This difference could be explained in at least two ways. Firstly, previous studies examining the positive effect of cognitive functions on physical activity relied on self-reported physical activity, which can bias the observed associations (Cheval, Orsholits, et al., 2020; Cheval et al., 2019; Lindwall et al., 2012). Secondly, our study relied on general cognitive functioning, whereas previous results highlight the specific importance of inhibition resources that may be required to counteract an automatic tendency for effort minimization (Cheval & Boisgontier, 2021; Cheval et al., 2021; Cheval, Daou, et al., 2020; Cheval, Radel, et al., 2018; Cheval et al., 2019; Cheval, Tipura, et al., 2018). Therefore, future studies should investigate the specific relationships between motor inhibition and physical activity when such data is available.

### Strengths and limitations

Among the strengths of the current study are the use of large-scale datasets, the reliance on instruments derived from objective measures of physical activity, and the application of a robust genetically informed method that can estimate causal effects. However, this study has several features that limit the conclusions that can be drawn. First, the measure of cognitive functioning spans multiple performance domains, which reduced the specificity of the cognitive functioning that was assessed. This feature limits our ability to evaluate the putative causal effects between specific cognitive functioning, such as motor inhibition, and physical activity. Second, MR analysis is designed to elucidate a life-long exposure effect on a life-long outcome (except in special cases when genetic factors have time-dependent effects), thus it is not suited to explore temporal aspects of these causal relationships. Third, 2-sample MR methods require that SNP effects on the exposure are homogeneous between the two samples. Here, because our two samples differ in age, we rely on the assumption that these genetic effects do not change depending on age. This assumption often turns out to be true, although there are rare exceptions (Winkler et al., 2015). It is therefore still possible that genetic variants related to physical activity and cognitive function may differ across the life course. For example, genetic variants related to cognitive development, maintenance and decline may strongly differ. Likewise, the genetic variance predicting physical engagement in early-life may differ from those predicting engagement in adult or late-life. Accordingly, as the age range between the sample is not equivalent (40 to 60 years for the UK Biobank vs 8 to 96 years in the COGENT consortium) and, most importantly, as physical activity was only assessed in the UK biobank that provides the narrowest age range, the potential differences in the genetic variants depending on individual’s age may have bias the current findings. Testing to which extent age may influence the genetic variants associated with physical activity and cognitive functioning traits is thus warranted in future studies. Fourth, LHC-MR can be limited by the low heritability of traits, potentially causing bimodal/unreliable estimates. Fifth, LHC-MR assumes a single confounder (or several ones with similar effects), but a limitation exists when multiple confounders are present with similar but opposing effect directions on the traits of interest, resulting in a higher misdetection rate. Sixth, alhough the coefficients estimated with LHC-MR did not statistical differed from the coefficents estimated with classical MR, it is important to akcnowledge that no classical MR were unable to find a significant association from physical activity to cognitive function. Accordongly, even if we can be rather confident in the estimation provided by the newly developed methods, it seems more reasonable to consider that the current findings are provisional and need to be replicated. Finally, it is worth noting that the genetic instruments were developed on a primarily white population of European ancestry, limiting the generelizeability of the results.

## Conclusion and policy implications

Our findings provide preliminary support for a unidirectional relation whereby higher levels of moderate and vigorous physical activity lead to improved cognitive functioning. These results underline the essential role of moderate and vigorous physical activity in maintaining or improving general cognitive functioning. Therefore, health policies and interventions that promote moderate and vigorous physical activity are relevant to improve cognitive functioning or to delay its decline.

## Acknowledgements

B.C. is supported by an Ambizione grant (PZ00P1_180040) from the Swiss National Science Foundation (SNSF). M.P.B. is supported by the Natural Sciences and Engineering Research Council of Canada (RGPIN-2021-03153) and the Banting Research Foundation. Y.C.K. is supported by the National Institute of Health (R01 HL136528). The authors are thankful to Gail Davies for providing useful comments on the manuscript.

